# Aberrant cocoons found on honey bee comb cells are found to be *Osmia cornifrons* (Radoszkowski) (Hymenoptera: Megachilidae)

**DOI:** 10.1101/2019.12.16.875856

**Authors:** Francisco Posada-Florez, Barbara Bloetscher, Dawn Lopez, Monica Pava-Ripoll, Curtis Rogers, Jay D. Evans

**Affiliations:** Bee Research Laboratory, Agricultural Research Service, UDSA, Beltsville, MD 20705; Ohio Dept of Agriculture, Division of Plant Health; U.S. Food and Drug Administration, Center for Food Safety and Applied Nutrition, College Park, Maryland, USA

**Keywords:** Solitary bee, nesting, supersedure, cleptobiosis, surrogate, inquiline, invasive species

## Abstract

Potential biological threats to honey bees must be addressed and validated quickly, before making disruptive and costly decisions. Here we describe numerous *Osmia cornifrons* (Hymenoptera: Megachilidae) cocoons in honey bee cells from one bee hive in Ohio. The developing *Osmia* cells presented themselves as a mystery at first, catching the attention of regulatory agencies. Along with identifying this species as a presumably benign resident in honey bee colonies, our observations suggest *Osmia* may use stored honey bee resources to provision offspring. Conceivably, resident honey bees might even act as surrogates, by provisioning *Osmia* offspring with pollen. Since the cocoons were attached to one another with honey bee wax, it seems likely that honey bee hosts were present during *Osmia* development. *Osmia* females have some plasticity when selecting nesting resources, and, upon discovering honey bee comb can use this resource for raising offspring. Along with resolving a potentially new biotic threat to honey bees, this diagnosis suggests a method for mass production of *Osmia* pollinators using an array of single cell foundation.

## Introduction

The U.S. beekeeping industry is constantly alert for the presence of organisms that may threaten or cause damage or losses to honey bee hives and the pollination services they provide. Examples of recently introduced threats to the honey bee industry include: the honey bee tracheal mite (*Acarapis woodi* (Rennie)), Small hive beetle (*Aethina tumida* Murray) and the Varroa mite (*Varroa destructor* (Anderson and Trueman)). There are no individual estimations of the costs of damages and loses, but these pests contribute directly or indirectly with colony losses, accounting for around 42.1% of mortality of colonies in the U.S. (Bee Informed Partnership, 2019).

Various invasive species of bees, wasps, flies, mites and other arthropods have the potential to threaten the honey bee industry in the U.S. They may cause direct damage or vector infectious diseases (Bram et al., 2002; Spivak et al., 2011; Core et al., 2012; Monceau et al., 2014; McMenamin & Flenniken, 2018). The introduction of a possible pest or invasive species that will compete for a niche, habitat or other resources is undesirable (Le conte & Navajas, 2008). In addition, enormous financial resources are needed for controlling new pests and parasites, along with training beekeepers on new pest management.

Recently, a sample of honey bee comb with cells containing unidentified cocoons was sent to the USDA-ARS Bee Research Laboratory (BRL) in Beltsville, MD, from the Ohio State Apiary Inspector, for analysis and identification. The main objective of the BRL was a rapid identification to ensure the species responsible for these cells did not present a risk to honey bees. In the end, we describe the use of a honey bee hive by the mason bee *Osmia cornifrons*. This species is not likely to cause a major threat to honey bees and indeed this result suggests potentially new ways to foster the success of solitary cavity-nesting bees.

## Materials and methods

### Record of the samples

The suspicious comb sample was received at the Diagnostic Lab of the BRL in Beltsville, MD in December of 2018. The sample was sent by the Honey Bee Inspector at the Ohio Department of Agriculture. The record stated the sample was taken in September, 2018, from an active honey bee colony housed within a standard Langstroth ten-frame honey bee hive. The honey bees in this hive were populated from a colony removed in April, 2018, from the wall of a cabin with Global Positioning System (GPS) coordinates of 40.1041 N −81.1630 W.

### Sample evaluation

The honey bee comb with the unidentified cocoons was analyzed visually to uncover clues for identifying potentially invasive organisms present in the comb sample. Individual cocoons were removed with sterile forceps from the honey bee comb to evaluate and document the characteristic shape, size and color. Photo documentation was undertaken to illustrate the process of identification, as this case was considered highly unusual.

### Morphological identification

First, the external characteristics of the cocoon surfaces were observed by stereomicroscope. The cocoons were also cut open to observe, describe, and identify the internal contents using morphological characteristics. These observations sought to explain any behavior or ecological interactions of these cocoons and the honey bee comb they were found in.

Samples of the organisms found inside the cocoons were placed in vials with 70% alcohol and labeled for later studies and comparisons. Photographs of all samples were taken using a Zeiss camera attached to a stereomicroscope for illustration and identification.

Alongside the examination of the samples, a literature search was done to locate records on this type of insect cocoon, bees cell capping, and modification of nest behavior of solitary bees.

### Molecular identification

DNA was extracted from one individual pre-pupa that was pulled out from a cocoon sample (voucher # USDA-BRL 181221-03_G21; Fig 1-9 to 1-13) using the DNeasy Blood & Tissue Kit (Qiagen Inc., Valencia, CA), as per manufacturers protocol, and eluting in 200 µl of molecular grade water. The cytochrome c oxidase subunit I (CO1) gene fragment was amplified using conventional PCR with primers LCO1490 GGTCAACAAATCATAAAGATATTGG and HCO2198 TAAACTTCAGGGTGACCAAAAAATCA (Folmer et al., 1994), with the following cycling parameters: 2 minutes at 95°C, followed by 30 cycles of 5s at 95°C, 45s at 50°C and 45s at 72°C and ending with a final phase of 72°C for 3 minutes. PCR products were visualized on a 1.5% agarose gel and cleaned using the HiPure PCR product Cleanup kit (Roche Life Sciences, Indianapolis, IN) and sent for sequencing at Macrogen USA (Rockville, MD) using LCO1490/HCO2198 sequencing primers. Simultaneously, one honey bee adult from another honey bee hive (voucher # USDA-BRL 181221-03 K21) was used as positive control through all steps starting with the DNA extraction for protocol validation. Sequences were compared against those in the NCBI nucleotide BLAST(nr/nt) database (http://blast.ncbi.nlm.nih.gov) using the megablast algorithm. Percent identities >99% were considered reliable identifications for this organism.

**Figure 1.**
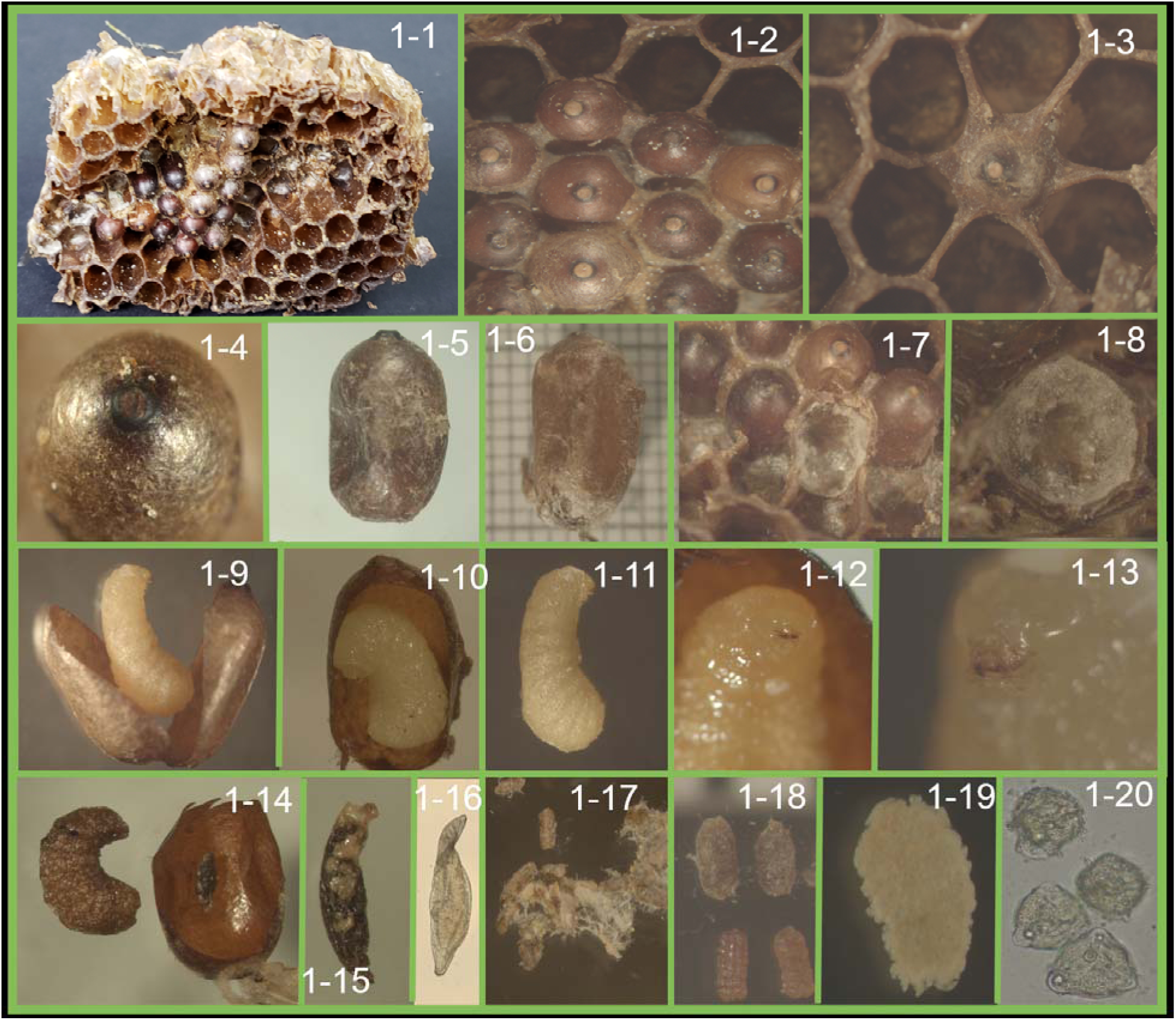
Diagnosis of aberrant cocoons found on honey bee comb cells. 1-1. Sample of honey bee comb with cells filled with cocoons. 1-2. Cocoons inserted in honey bee cells and attached to it with honey bee wax with a smooth form. 1.3. Cocoons located and surrounded by honey bee cells.1-4. Superficial view of cocoon showing a nipple formation. 1-5. Vertical view of the cocoon, showing a nipple formation and cylindrical shape.1-6. Measurement of the cocoon with attached pieces of silk. 1-7. Shows how the cocoons are inserted deeply in the comb cells and adhered to the walls of the cell with silk and wax. 1-8. Bottom of honey bee cell showing a cushion of silk where *O. cornifrons* larva deposits. 1-9 and 1-10. Cocoon cut open revealing the larvae. 1-11. Larva typical of vermiform hymenoptera apocrita. 1-12 and 1-13. Larva showing the head and mouth parts with mandibles. 1-14. Cocoon with larvae of O. *cornifrons* showing a dark spot on the fourth segment, a sign of parasitism by a wasp that remains on the cocoon. 1-15. Pre-pupa of the parasitoid wasp. 1-16. Egg of the parasitoid was found on the cocoon remains.1-17. Feces of *O. cornifrons* wrapped in silk. 1-18. Feces of *O. cornifrons* (top) and wax moth (bottom). 1-19, Feces of *O. cornifrons* larvae that aggregated when wet and placed under the microscope slide.1-20. Pollen grain observed on the *O. cornifrons* feces.

Sanger-derived nucleotide sequence for voucher # USDA-BRL 181221-03_G21 was aligned using CLUSTALW of the Molecular Evolutionary Genetics Analysis (MEGA) version X (Kumar et al., 2018) against 25 CO1 sequences retrieved form GenBank belonging to bees in the genus *Osmia*, the family Megachilidae, and the Apoidea bees as a whole. A Maximum Likelihood (ML) phylogenetic tree with 500 bootstraps was generated using default parameters in MEGA version X (Kumar et al., 2018), with the honey bee, *Apis mellifera* voucher # USDA-BRL 181221-03 K21 as an outgroup.

## Results

### Sample evaluation

The cluster of unusual cells attached to and embedded in the honey bee comb cells was peculiar (Figure 1-1). The sample comb was consistent with that of *A. mellifera* made of wax, hexagonal in shape and with a size of 5.1 ± 0.01 millimeter (mean ± SD, n=10). No honey bee life stages were present in the comb. Debris and feces of wax moth were present, indicating moths were feeding on the comb.

Clusters of cocoons were on both sides of the frame, surrounded by empty honey bee cells, and in the same general location on both sides (side one n=30 and side two n=24 cocoons). They were placed in a horizontal position inside the honey bee cells, and built in the comb as the honey bee would (Figure 1-1). The cocoons were inserted and well attached in the cells with bee wax surrounding the cocoon and just the apical tip of the cocoon exposed (Figure 1-2). The color of the cocoons was brown with a striking terminal cap having a hard black spot resembling a “nipple” (Figure 1-3 to 1-6). The cocoon shape was cylindrical with a longitudinal size of 10.0 ± 1.1 millimeters and an equatorial size of 6.0 ± 0.5 millimeters (Figure 1-5 and 1-6). Externally, the cocoons were wrapped in a thin layer of brown silk and were spun of a leathery substrate, hard to cut and water resistant (Figure 1-5, 1-6, 1-9, 1-10).

On the bottom of the honey bee cells, where the cocoons were attached, was a cushion of silk and the remains of pollen with feces, resembling sausages wrapped with silk. The analysis of this feces showed that the content was digested pollen (Figure 1-7, 1-8, 1-17 to 1-20) not belonging to honey bees, or other arthropods that typically thrive in honey bee colonies.

### Morphological identification

The first visual examination of the sample and corresponding internet and publication searches for identification gave close visual matches to drone brood cells of *Apis cerana* (Hymenoptera: Apidae), complete with a domed capping and a nipple formation. However, upon closer inspection, the building material of the cocoons themselves was not wax, even though the borders connecting the cocoons were attached with honey bee wax. Several cocoons were cut open to expose the larvae or yellowish pre-pupae. The shape of the larvae was more consistent with Hymenoptera than Diptera. Observing the setal process and chaetotaxy of the larvae, it revealed they were smooth with few segmentations and the head showed a mandible structure that resembled most closely aculeate Hymenoptera (Figure 1-9 to 1-13).

The combined features of cocoons are of reddish-brown color, wrapped with a thin layer of silk in their exterior, tough skin, and with a nipple formation on the protruding cap (Figure 1-4 to 1-6). These features suggest these were cocoons of the Mason bee (Hymenoptera: Apoidae: Megachilidae) that were deposited on the honey bee comb.

After cutting open the cocoons, 30 larvae were found as pre-pupa, with different buccal morphologies (Figure 1-11 to 1-13). Interestingly, a parasitoid wasp was found inside one cocoon, but external of the larva, indicating this species is an ectoparasitoid of larvae. The egg’s corium was also found inside the cocoon, confirming that after the egg hatched, the larva of the parasitoid moved to the *O. cornifrons* larva to feed externally. The examination of the wasp morphology suggests it belongs to the superfamily Chalcidoidea.

### Molecular identification

The molecular identification of the pre-pupa sample (voucher # USDA-BRL 181221-03_G21), was confirmed to be the Japanese hornfaced bee, *Osmia cornifrons* Radoszkowski (Hymenoptera: Apoidea: Megachilidae). BLASTN results of the sequenced COI fragment (686 bp) had 100% identity to nucleotide sequence of another *O. cornifrons* isolate deposited in GenBank (EU726549). Additionally, the maximum likelihood (ML) phylogenetic tree constructed with other Megachilidae and Apoidae species showed a good bootstrap support for the sample *O. cornifrons* isolates (Figure 2). CO1 nucleotide sequences for *O. cornifrons* voucher # USDA-BRL 181221-03_G21 and *A. mellifera* voucher # USDA-BRL 181221-03 K21 were deposited in GenBank under accession numbers MN091623 and MN091624, respectively.

**Figure 2.**
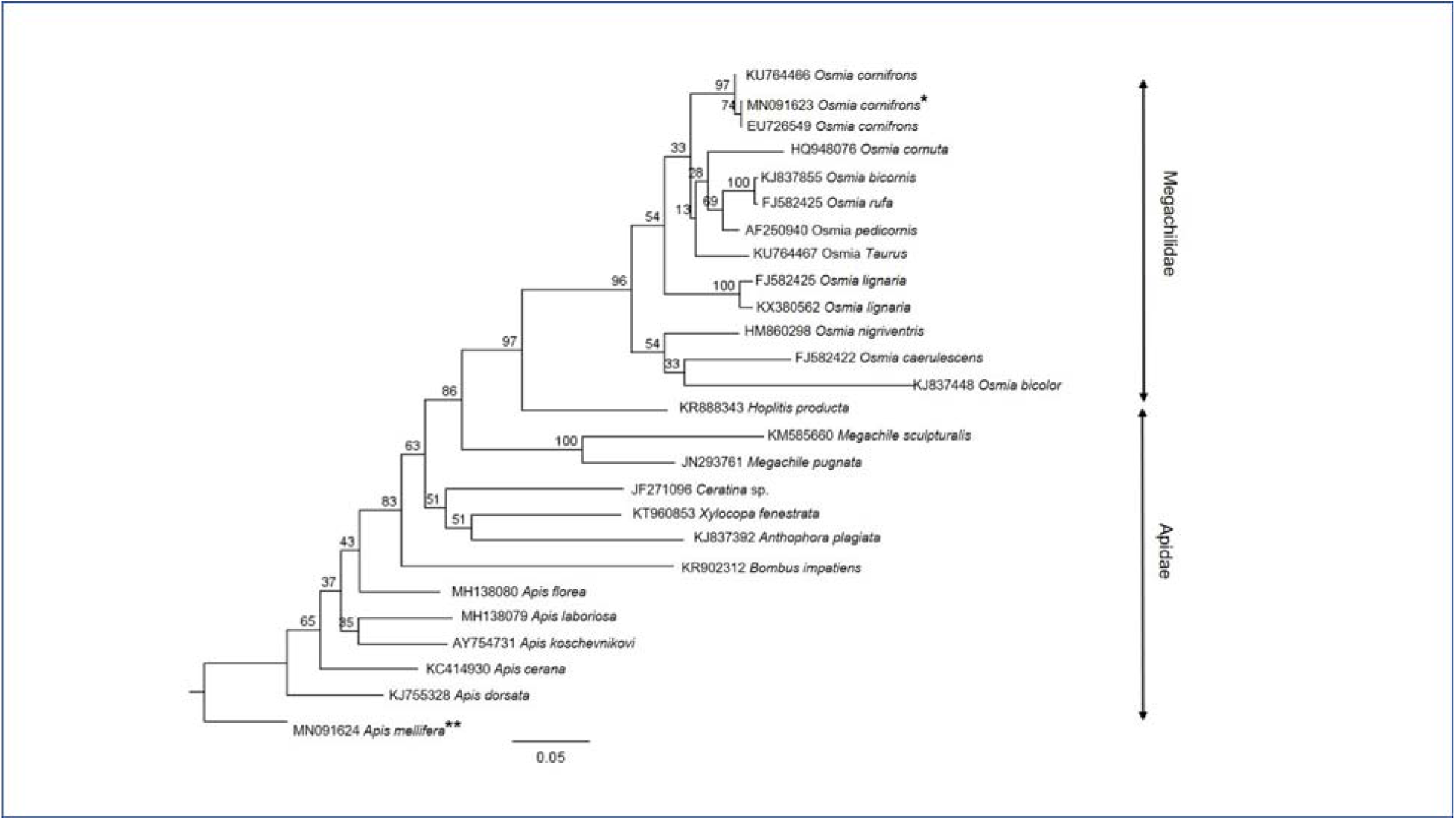
Phylogenetic tree of cytochrome c oxidase subunit 1 (CO1) nucleotide sequences of *O. cornifrons* GenBank accession # MN091623 (*voucher # USDA-BRL 181221-03_G21) and other representatives from Megachilidae and Apidae families, including *A. mellifera* GenBank accession # MN091624 (**voucher # USDA-BRL 181221-03 K21). The tree was constructed based on the alignment of the CO1 sequences using MEGA X and the Maximum Likelihood method. Numbers on branches indicate the bootstrap values obtained after 500 replications.

## Discussion

*Apis cerana*, and its associated pests, are not found in North America and their possible presence in honey comb raised alarm because this is a reportable species by USDA Animal Plant Health Inspection Service (APHIS). Bees of the genus *Apis* use wax to build the cells and cap their brood. However, the building material of the examined cocoons was not found to be wax, therefore cocoons reported here did not belong to the genus *Apis*. Similarly, the cells did not match those vespid wasps that use paper to build the cells and silk to cap them. Neither belong to sphecid wasps that use mud to build entire cell. None of these materials were found in the examined cocoons.

The cocoons had a superficial resemblance to fly puparia. Conopidae flies are reported to parasitize honey bees and other species of bees (Abdalla et al., 2014; Gibson et al., 2014). However, these flies are endoparasites and the puparium usually has two terminal protuberances (Abdalla et al., 2014). Also, conopids pupate individually in the ground and there are no records of gregarious pupation behavior. Calliphoridae flies also can attack honey bees (Morse & Flottum, 1997), but it is unusual for so many cocoons to be produced in a comb in such an organized fashion (Figure 1-1 to 1-3). Besides flies, other cocoons with features similar to the cocoons in the sample are Braconid wasps. However, Braconid wasp cocoons are made of silk and the examined cocoons only had a thin layer of silk externally and were larger than Braconid wasp cocoons.

All findings described in the results section suggest the cocoons in the sample belong to the Mason bee, *Osmia cornifrons*. Mason bees generally nest in cavities with several offspring lined up end-to-end (Bosch et al., 2008; Boyle & Pitts-Singer, 2019) and show a preference for twigs or tubes, like bamboo, or man-made nesting tubes or tunnels. They are solitary when building their nests, but can be gregarious. Female bees pack cells with pollen and enclose the food within the cell after they lay the eggs, then seal the cavities with mud (Cane et al., 2007; Mckinney, 2011). However, none of these features were present on the honey bee comb examined, except that the honey bee comb cells resemble the holes that Mason bees nest in. We can speculate that one or more *O. cornifrons* females laid eggs in expanded honey bee comb cells after provisioning these cells with pollen. The fact that the cocoons were attached to cells with honey bee wax suggests that honey bees were present during development (Figure 1-2 and 1-3). One possibility is that *O. cornifrons* also exploit food resources within honey bee hives, or even that honey bees themselves took part in provisioning, although there were no observations to demonstrate this.

The life cycle of *O. cornifrons* states that the larval stage occurs from May-June to September-October. This is consistent with the phenology found for this species (Bosch et al., 2008; Mckinney, 2011; Boyle & Pitts-Singer, 2019) and with our collection of these samples in September. Nesting of *O. cornifrons* within honey bee hives did not seem to have any effect on the developmental rate of the life cycle. Additionally, one of us (BB) put combs containing these cells in an incubator for several weeks to try and rear adult insects to emerge for identification, to no avail. This is also consistent for *O. cornifrons*, as they would not normally emerge until spring. The Chalcidoidea wasp found inside the cocoons has been also reported as parasitoids of solitary bees (Figure 1-14 to 1-16) (Krunic et al., 2005).

The Megachilidae family, to which the *O. cornifrons* belongs, has some species that are inquilines in cells of other family members (Clausen, 1940), but *O. cornifrons* is not known to do so. Megachilidae bees can be raised for pollination services for orchard crops and seed production (Bohart, 1972; Cane et al., 2007). Several records have shown that Mason bees nest in different structures, such as abandoned snail shells and in sphecid wasp nests, including one supersedure event which destroyed the nest of *Isodontia mexicana* (Saussure) (Hymenoptera: Sphecidae) (Delphia & O’neill, 2012). Supersedure has also been reported between species of *Osmia*, possibly competing for nesting sites (Bohart, 1955). *Osmia inermis* (Zetterstedt) was found forming clusters inside flat containers when they were provided as artificial nesting (Sheffield et al., 2015). Here, we report the first record of *O. cornifrons* nesting on combs of honey bee cells, further demonstrating its nesting plasticity. Another interpretation of this changing nesting behavior could be due to habitat transformation that put pressure on the species to find adequate places for nesting (Sheffield et al., 2015).

This information demonstrates that *O. cornifrons* continues its establishment and distribution into the West since its introduction from Japan in 1977 - 1978 to Beltsville, MD as a beneficial insect for pollination (Batra, 1978). Before being introduced, *O. cornifrons* was placed into quarantine to avoid the introduction of undesirable parasitoids, predators and diseases. A possible application of these findings is to further research the use of templates, resembling the honey bee comb, built with materials that allow the harvesting of cocoons for commercial purposes. If bees could be lured to nest in single-bee cells, this could improve protection against natural enemies and possibly prevent the detrimental impact from parasitic mites, parasitoids and microbes that attack *Osmia* spp. (Krunic et al., 2005; Park et al., 2009). With this combined information, we now must try to understand how the Japanese hornfaced bee, *O. cornifrons*, used honey bee cells as a nesting site. Additional questions include: Are these bees changing their nesting behavior? How could they use the space on the comb? Did they take advantage of stored honey bee pollen? Why were they not attacked and destroyed by honey bee workers that meticulously guard and defend the hive from such intruders?

In summary, the evidence here suggests that honey bees took care of, or at least tolerated, larvae of *O. cornifrons* that developed inside cocoons attached with honey bee wax (Figure 1-1 to 1-3). These cocoons were built by the silk produced by larvae combined with a hardened secretion, like leather (Figure 1-5 and 1-6). The cocoons resemble a coarctate pupa and the larval shape was typical for Hymenoptera, with a distinctive head and mouth parts (Triplehorn & Johnson, 2005) (Figure 1-9 to 1-13).

## Acknowledgments

The authors would like to give special thanks to Joe Heider, the beekeeper who found *O. cornifrons* cocoons nesting in honey bee hives and who sent them to Barbara Bloetscher. Francisco Posada-Flórez would like to express his gratitude to the ORAU/ORISE fellowship program awarded through USDA-ARS.

